# Human-dose equivalent 5-fluorouracil triggers a pathophysiological cascade of neuroinflammation, cortical remodeling, and behavioural disruption in Wistar rats

**DOI:** 10.1101/2025.09.15.676357

**Authors:** Anthony T. Olofinnade, Ajao Joshua Aduramigba, Akinsehinwa Akinsanmi Festus, Olakunle J. Onaolapo, Adejoke Y. Onaolapo

**Author notes:** **Corresponding author Dr. (Mrs.) Adejoke Y. Onaolapo**, Behavioural Neuroscience/Neurobiology Unit, Department of Anatomy, Ladoke Akintola University of Technology, Ogbomosho, Oyo State, Nigeria.

## Abstract

5-Fluorouracil (5-FU) is an antimetabolite widely used in cancer chemotherapy. Despite its central role in many anticancer regimens information on its potential effects on the central nervous system (CNS) remains limited. This study investigated the effects of human dose equivalent 5-FU on the cerebral cortex of rats. Fifty male rats were randomly assigned into five groups (n=10). The control group received intraperitoneal (i.p.) saline, while four experimental groups received i.p. 5-FU at 12.5 25 50 or 100 mg/kg body weight. Saline or 5-FU was administered for four consecutive days, followed by alternate-day dosing until day 12. Behavioral assessments were conducted and animals were sacrificed 24 hours after the final test. Cortical tissues were analyzed using biochemical assays histology and immunohistochemistry. 5-FU administration caused dose-dependent decreases in body weight food intake and locomotor activity. Treated rats also showed impaired spatial working memory and reduced time spent in the open arms of the elevated plus maze. Biochemically 5-FU significantly increased cortical malondialdehyde and tumor necrosis factor-alpha levels while total antioxidant capacity and interleukin-10 levels decreased. Histological analysis revealed progressive disruption of cortical cytoarchitecture with increasing doses. 5-FU induces dose-dependent neurotoxicity in the rat cerebral cortex characterized by behavioral deficits, oxidative stress neuroinflammation and histomorphological alterations. These findings highlight the need to better understand and mitigate CNS toxicity associated with 5-FU–based chemotherapy.

## 1. Introduction

Fluoropyrimidines, including floxuridine, capecitabine, and fluorouracil (5-FU), are antimetabolite chemotherapeutic agents that have remained central to the management of solid organ cancers for several decades [1,2]. Both monotherapy and combination regimens incorporating fluoropyrimidines continue to be standard in the treatment of metastatic disease and in adjuvant protocols [2]. Among them, 5-FU, first developed in the 1950s, is one of the most widely prescribed agents for gastrointestinal, breast, and other solid tumors [1].

The cytotoxic effects of 5-FU are mediated primarily through inhibition of thymidylate synthase, which disrupts DNA synthesis and replication, as well as through incorporation of its metabolites into RNA and DNA [1,4–7]. Clinically, intravenous 5-FU is approved for gastric, breast, pancreatic, and colorectal adenocarcinomas, while topical formulations are employed for premalignant and malignant skin conditions such as actinic keratoses and superficial basal cell carcinomas [3]. Despite these therapeutic benefits, 5-FU is associated with systemic toxicities that limit its clinical use.

Beyond peripheral toxicity, chemotherapeutic agents, including 5-FU have been increasingly recognized to exert adverse effects on the central nervous system (CNS). These effects manifest as deficits in attention, memory, and executive function, collectively termed “chemo-brain” or “chemo-fog” [8–11]. While often considered nonspecific consequences of treatment, persistent cognitive impairment suggests that certain agents may induce specific neurotoxic changes [10,11]. Indeed, although less frequently reported, 5-FU has been implicated in neurological syndromes such as multifocal leukoencephalopathy [12] and Wernicke’s encephalopathy [13]. Experimental studies have further demonstrated that 5-FU can induce anxiety-like behavior and impair spatial memory in rodents [14, 15].

Despite its longstanding use and widespread clinical relevance, the neurotoxic profile of 5-FU particularly its impact on cortical integrity and function remains poorly characterized. The cerebral cortex, being integral to higher-order cognitive processes, represents a critical but understudied target of 5-FU toxicity. This study therefore seeks to examine the effects of 5-FU on the cerebral cortex of rats, with the aim of elucidating neurobehavioral, neurochemical, and structural alterations. Findings from this work may advance understanding of chemotherapy-associated cognitive dysfunction and provide insights into mechanisms underlying 5-FU– induced CNS toxicity.

## 2. Methods

### 2.1 Chemicals and drugs

5-fluorouracil (5 Flucel^®^, 250 mg/5 ml injection), Normal Saline, Assay kits

### 2.2 Animals

Animals used in this study were sourced from the animal house of the Ladoke Akintola University of Technology Ogbomoso, Oyo State, Nigeria. Rats were housed in wooden cages that measured 20 x 10 x 12 inches in temperature-controlled (22.5°C ±2.5°C) quarters with lights on at 7.00 am. Rats were allowed free access to food and water. All procedures were conducted in accordance with the approved protocols of the Faculty of Basic Medical Sciences, Ladoke Akintola University of Technology and within the provisions for animal care and use according to the European Council Directive, (EU2010/63) on scientific procedures in living animals.

### 2.3 Diet

All animals were fed commercially-available chow (29% protein, 58% carbohydrate. 11% fat) throughout the study.

### 2.4 Experimental methodology

Fifty adult male rats weighing 150-170g each were assigned into five groups of ten (n-10) rats each. The groups included vehicle control that were administered intraperitoneal (i.p.) injection of saline at 2 ml/kg, and four other groups administered i.p. 5-Fluoruracil at 12.5, 25, 50 and 100 mg/kg body weight respectively. Saline and 5-Fluorouracil were administered for four consecutive days initially, then on alternate days 6, 8, 10 and 12 [16]. The doses of 5-Fu translate to human equivalent doses that are within the clinical dose range [14]. At the end of the experimental period, animals were exposed to the open-field arena (for the assessment of locomotor, rearing and grooming) behaviours, the elevated plus maze (EPM) for anxiety related behaviours, and the Y-maze for the assessment of spatial working-memory. Twenty-four hours after the last behavioural test, animals were sacrificed. The brains of the rats were excised, sections of the cerebral cortex were homogenised (for the estimation of oxidative stress, antioxidant status, and inflammatory markers); or processed for paraffin-embedding, and stained with haematoxylin/eosin and cresyl violet for general histological study. Immunohistochemical staining was performed for neuron-specific enolase.

### 2.5 Estimation of body weight and feed intake

Estimation of body weight of animals in all groups was done weekly, while the estimation of daily feed consumption was made an electronic weighing scale (Mettler Toledo Type BD6000, Switzerland) as previously described [17, 18]. The percentage change in body weight or food intake for each animal was calculated using the following equation, and the results for all animals were computed to find the statistical mean:

### 2.6 Behavioural tests

#### 2.6.1 Open-field behaviours

The open-field arena in rodents is used to assess central effects of a drug or chemical compound as measured by exploratory activity, rearing and grooming behaviours. Locomotion, rearing and self-grooming behaviours were scored during ten minutes of exposure to the arena. The open-field arena is a rectangular-shaped box with a hard floor that measures 36 x 36 x 26 cm; and is made of white-painted wood. Its floor is divided into 16 equal squares with a permanent red marker. Placement of animals and scoring of the various parameters are as previously described [19, 20].

#### 2.6.2 Y-Maze Test

This Y-maze spatial working-memory test is based on the natural propensity of rodents to explore novel environments. The Y-maze paradigm is made of wood and has three identical arms arranged in the shape of a “Y.” Each arm is usually 15 inches long and 3.5 inches wide, with 3-inch-high walls. Each rat was placed in one of the arms of the Y maze and allowed to move freely within the compartments until its tail completely enters another arm. The sequence of arm entries was recorded as described previously [21, 22].

#### 2.6.3 Elevated plus Maze

The elevated plus maze (EPM) test was developed to examine the possible anxiolytic effects of drugs in rodents. The paradigm is based on the propensity of rodents for closed spaces. It is a cross-shaped platform with two open (50 x 10 x 0 cm) and two closed arms (50 x 10 x 30 cm) placed 50–70 cm above the ground. Placement of animals and scoring of the various parameters are as previously described [23, 24].

### 2.7 Brain Homogenisation

The cerebral cortex was homogenised (1:10, w/v) in ice-cold phosphate-buffered saline using a Teflon-glass homogeniser as previously described [25]. The homogenate was then centrifuged for 15 minutes at 5,000 revolutions/minute at 4 °C. The supernatant was immediately used to assay respective markers.

### 2.8 Biochemical Test

#### 2.8.1 Estimation of Lipid peroxidation

Lipid peroxidation levels were measured as malondialdehyde content as previously described [26, 27] Change in colour was measured using a spectrophotometer at 532 nm.

#### 2.8.2 Antioxidant activity

Total antioxidant capacity was determined using commercially available assay kit. Colour changes were measured as described previously described [28, 29].

#### 2.8.3 Tumour necrosis factor-α, Interleukin (IL)-10 and interleukin-6

Tumour necrosis factor-α and interleukin (IL)-10 were measured using enzyme-linked immunosorbent assay (ELISA) techniques with commercially available kits (Enzo Life Sciences Inc. NY, USA) designed to measure the ‘total’ (bound and unbound) amount of the respective cytokines as previously described [30. 31]. Interleukin 6 level was assayed using enzyme-linked immunosorbent assay (ELISA) techniques with commercially available kits (Enzo Life Sciences Inc. NY, USA) as previously described [32, 33].

### 2.9 Protocol for neuron specific enolase (NSE) Immunohistochemistry

Immunohistochemical test for neuron specific enolase (NSE) was performed using the NovocastraTM and NovolinkDM polymer detection system (Leica Biosystems, UK) and appropriate primary monoclonal antibodies as previously described [34, 35].

### 2.10 Photomicrography

Sections of the cerebral cortex were examined using a Sellon-Olympus trinocular microscope (XSZ-107E, China) with a digital camera (Canon Powershot 2500) attached, and photomicrographs taken.

### 2.11 Statistical analysis

Data was analysed using Chris Rorden’s ezANOVA for windows. One-way ANOVA was used to test the hypothesis. Tukey (HSD) test was used for Posthoc analysis. Results were expressed as mean ± S.E.M, and p< 0.001 considered significant.

## 3.0 Results

### 3.1 Effect of 5-fluorouracil on body weight and feed intake

Figure 1 shows the effect of 5-fluorouracil on mean relative change in body weight (upper panel) and mean relative change in feed intake (lower panel). Mean relative body weight decreased significantly (p < 0.001) with 5-FU at 12.5, 25, 50 and 100 mg/kg body weight compared to control. Mean relative feed intake also decreased significantly (p < 0.001) with 5-FU at 12.5, 25, 50 and 100 mg/kg body weight compared to control.

**Figure 1:**
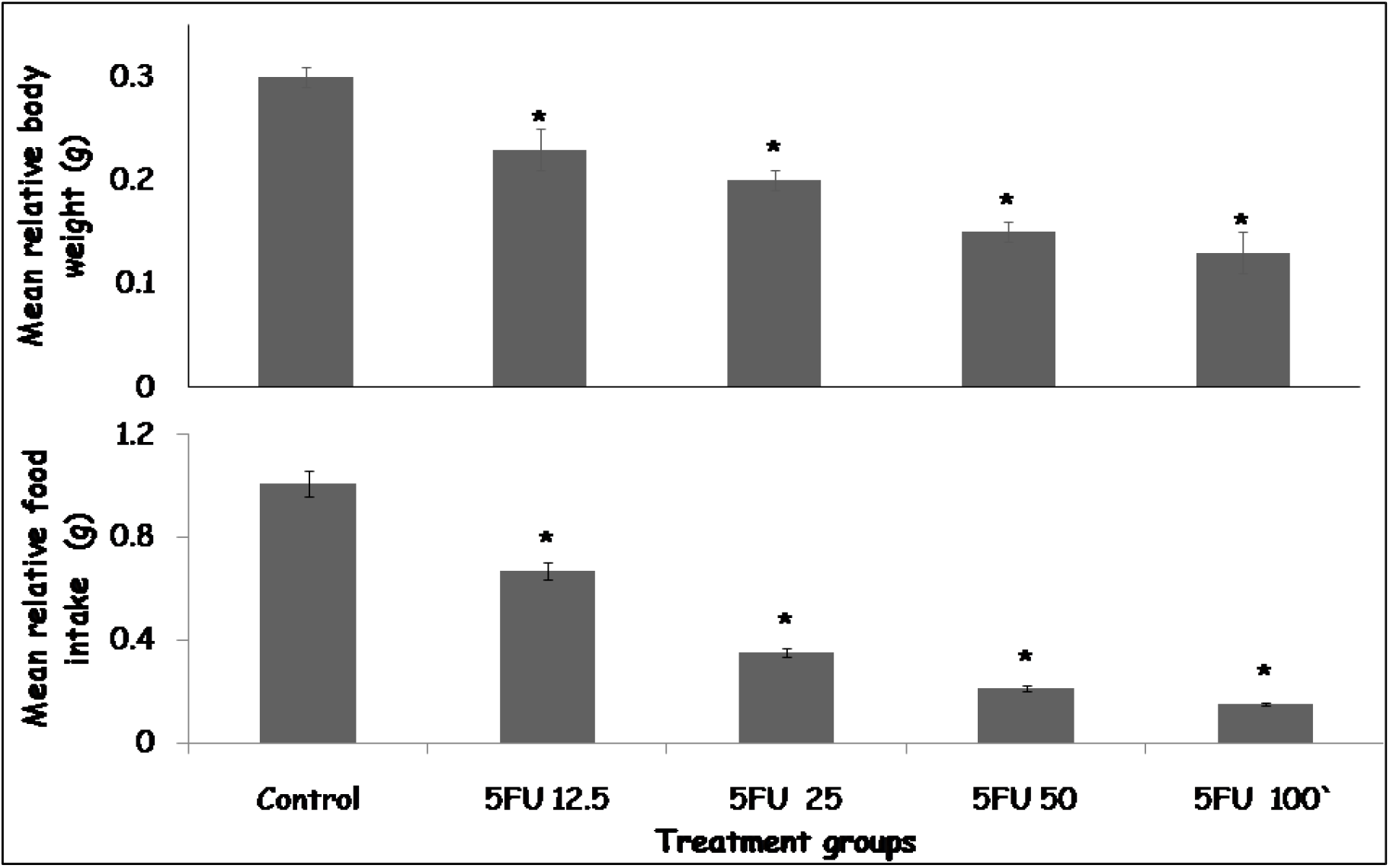
Effect 5-Fluorouracil on mean relative change in body weight (upper panel) and mean relative change in food intake (lower panel) in rats. Each bar represents Mean ± S.E.M, *p < 0.05 significant difference from control, number of rats per treatment group =10. 5FU: 5-Fluorouracil

### 3.2 Effect of 5-fluorouracil on exploratory behaviours in the open-field Arena

Figure 2 shows the effect of 5-fluorouracil on exploratory behaviours in the open field box scored as line crossing (upper panel) and rearing activity (lower panel). Line crossing decreased significantly (p < 0.001) with 5-FU at 12.5, 25, 50 and 100 mg/kg body weight compared to control. Rearing activity decreased significantly (p < 0.001) with 5-FU at 12.5, 25, 50 and 100 mg/kg body weight compared to control.

**Figure 2:**
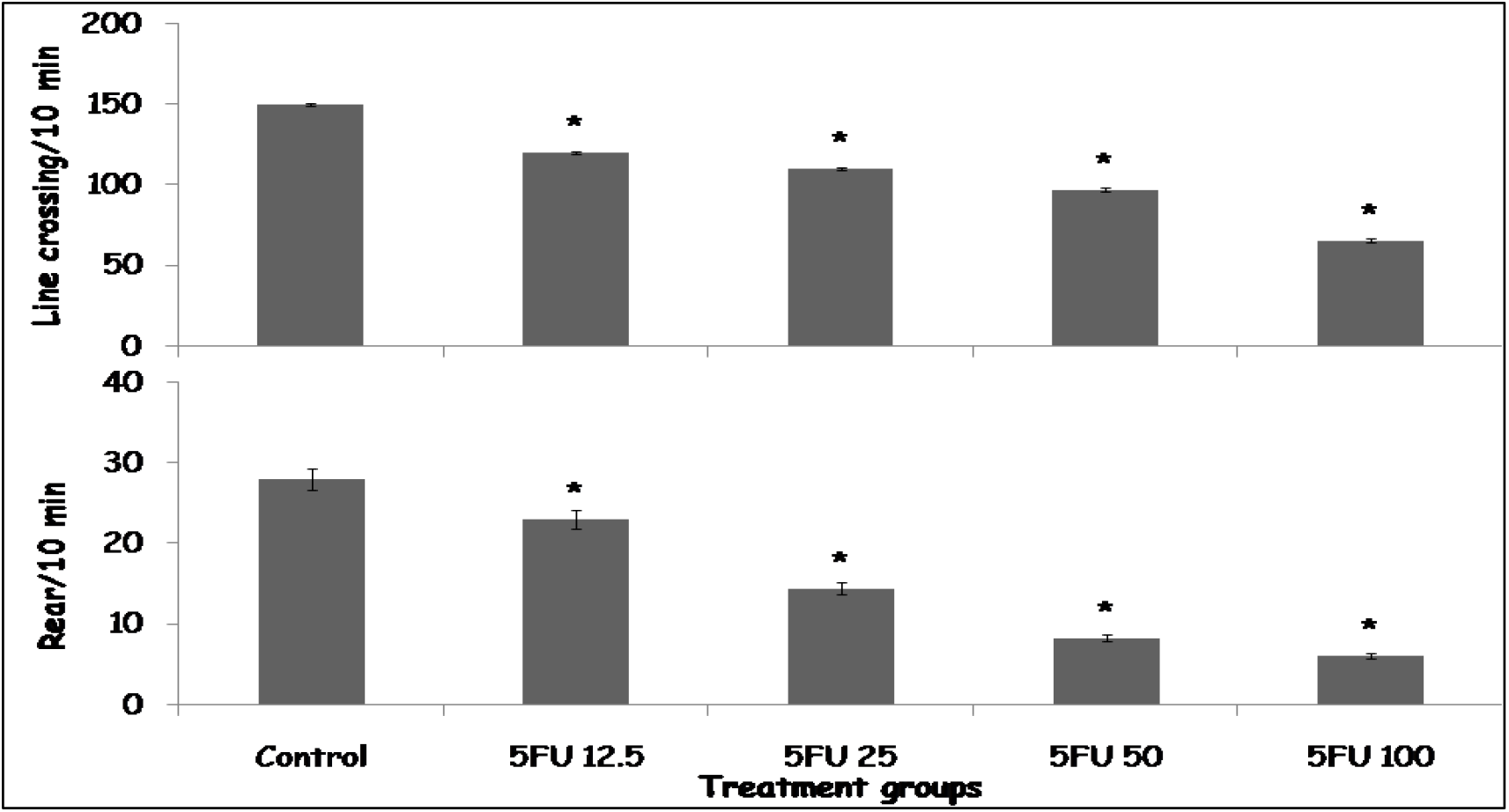
Effect 5-Fluorouracil on line crossing (upper panel) and rearing activity (lower panel) in rats. Each bar represents Mean ± S.E.M, *p < 0.05 significant difference from control, number of rats per treatment group =10. 5FU: 5-Fluorouracil.

### 3.3 Effect of 5-fluorouracil on Self-grooming in the open-field box

Figure 3 shows the effect of 5-fluorouracil on self-grooming behaviours in the open field box. Self-grooming decreased significantly (p < 0.001) with 5-FU at 12.5, 25, 50 and 100 mg/kg body weight compared to control.

**Figure 3:**
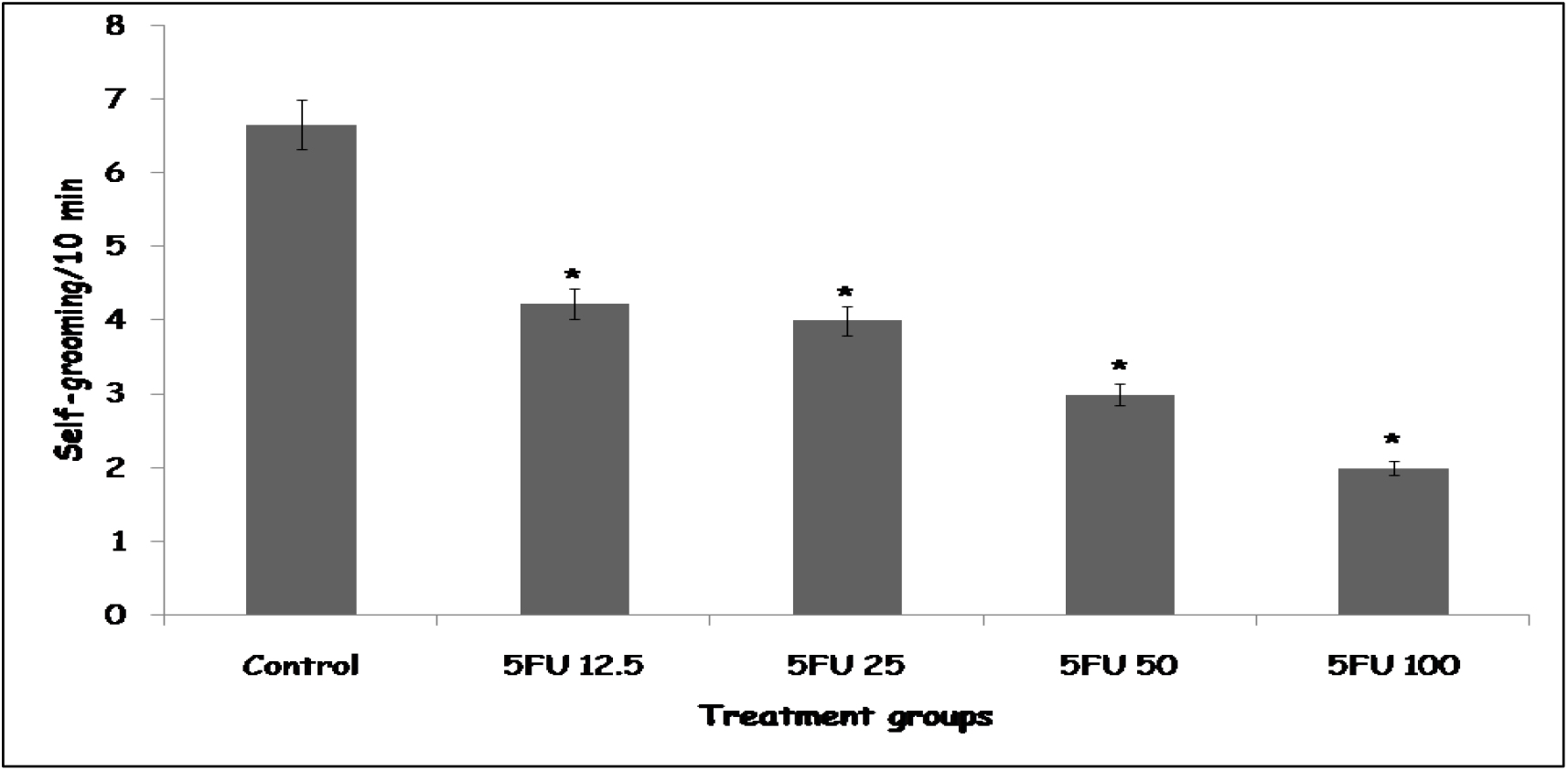
Effect 5-Fluorouracil on self-grooming behaviours in rats. Each bar represents Mean ± S.E.M, *p < 0.05 significant difference from control, number of rats per treatment group =10. 5FU: 5-Fluorouracil.

### 3.4 Effect of 5-fluorouracil on anxiety-related behaviours in the Elevated plus maze

Figure 4 shows the effect of 5-fluorouracil on anxiety-related behaviours in the elevated plus maze scored as time spent in the open arm (upper panel) and closed arm (lower panel). Time spent in the open arm decreased significantly (p < 0.001) with 5-FU at 12.5, 25, 50 and 100 mg/kg body weight compared to control. Time spent in the closed arm increased significantly (p < 0.001) with 5-FU at 12.5, 25, 50 and 100 mg/kg body weight compared to control.

**Figure 4:**
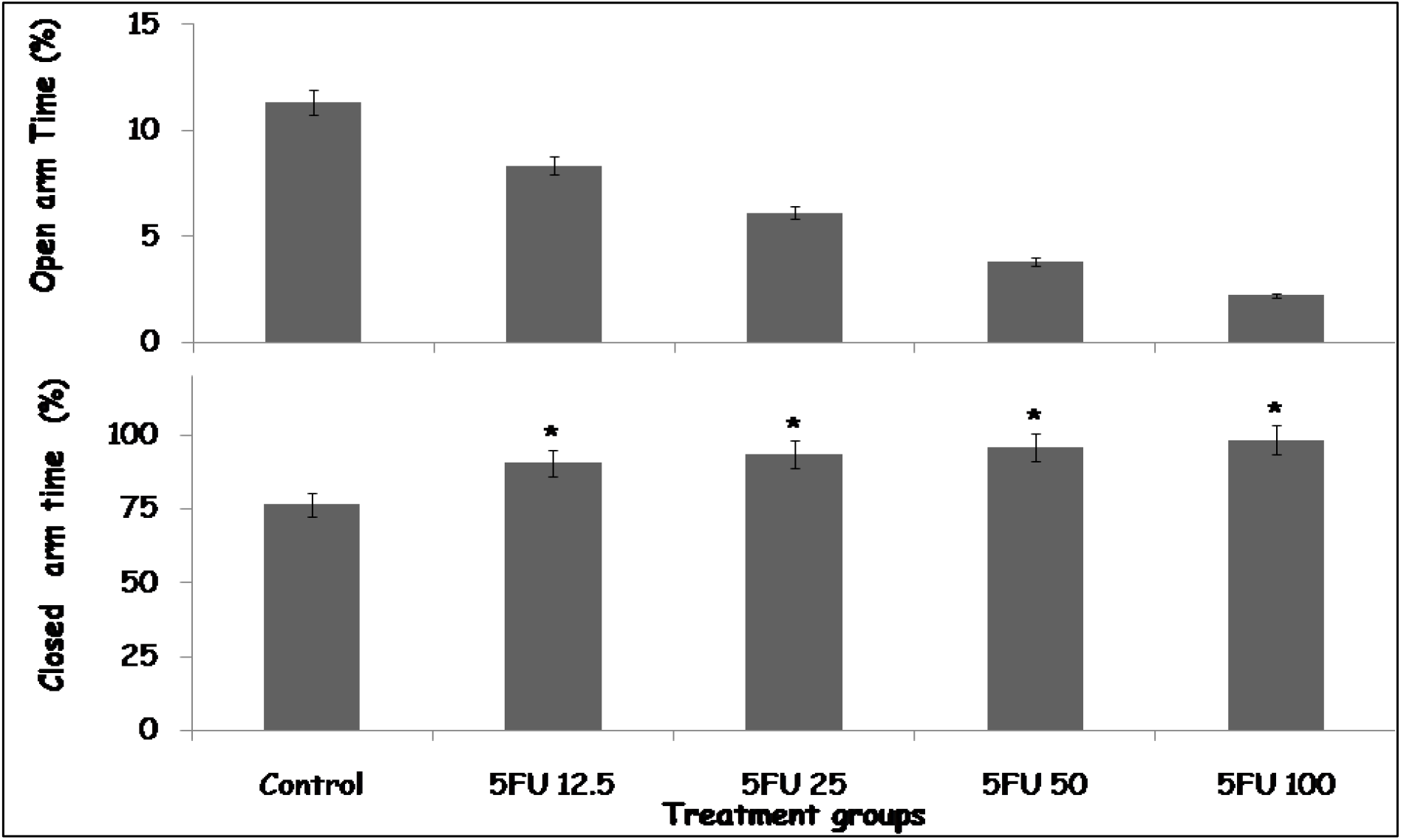
Effect 5-Fluorouracil on time spent in the open arm (upper panel) and closed arm (lower panel) of the elevated plus maze in rats. Each bar represents Mean ± S.E.M, *p < 0.05 significant difference from control, number of rats per treatment group =10. 5FU: 5-Fluorouracil

### 3.5 Effect of 5-fluorouracil on spatial working memory tasks in the Y and Radial arm maze

Figure 5 shows the effect of 5-fluorouracil on spatial working memory tasks in the Radial arm (upper panel) maze scored as alternation index and Y maze (lower panel) scored as percentage alternation in 5 minutes. Alternation index in the radial arm maze and percentage alternation in the Y maze decreased significantly (p < 0.001) with 5-FU at 12.5, 25, 50 and 100 mg/kg body weight compared to control.

**Figure 5:**
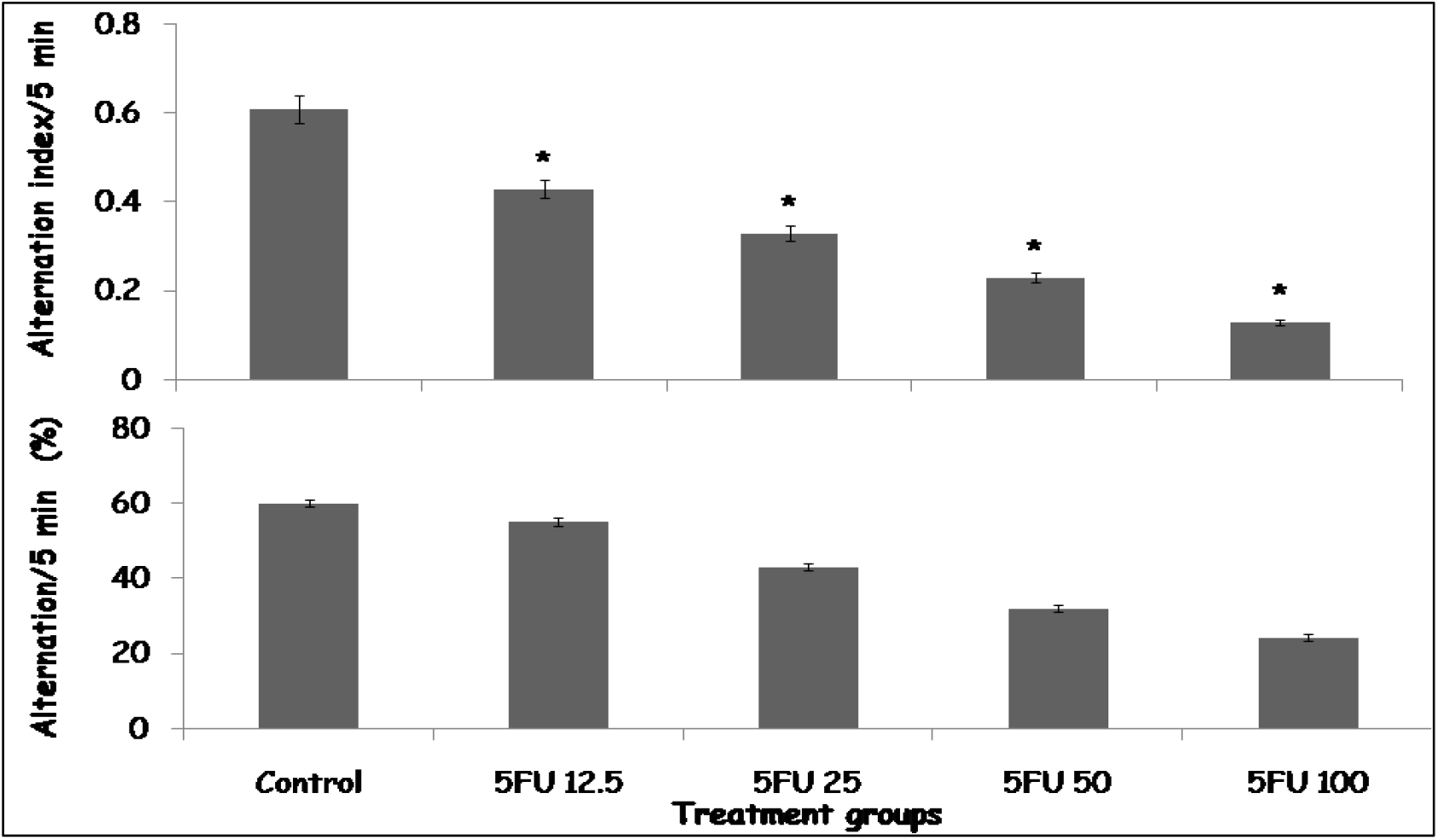
Effect 5-Fluorouracil on alternation index in the radial arm maze (upper panel) and percentage alternation in the Y maze (lower panel). Each bar represents Mean ± S.E.M, *p < 0.05 significant difference from control, number of rats per treatment group =10. 5FU: 5-Fluorouracil.

### 3.6 Effect of 5-fluorouracil on levels of lipid peroxidation, antioxidant capacity and inflammatory markers in the cerebral cortex

Table 1 shows the effect of 5-fluorouracil on lipid peroxidation, total antioxidant capacity, and inflammatory marker levels in the cerebral cortex. Lipid peroxidation measured as Malondialdehyde (MDA), Tumour necrosis factor (TNF)-α and Interleukin-6 levels increased significantly (p < 0.001) with 5FU at 12.5, 25, 50 and 100 mg/kg body weight compared to control. Total antioxidant capacity (TAC) and interleukin 10 levels decreased significantly (p < 0.001) with 5FU at 12.5, 25, 50 and 100 mg/kg body weight compared to control

**Table 1:** Effect of 5-fluorouracil on lipid peroxidation, antioxidant capacity and inflammatory markers.

| Groups | MDA (μm) | TAC (mM TE) | TNF-α ng/L | IL-10 pg/ml | IL-6 pg/ml |
| --- | --- | --- | --- | --- | --- |
| Control | 2.23±0.12 | 12.34 ±0.11 | 25.23±1.11 | 30.22±0.10 | 5.57 ± 1.23 |
| 5FU 12.5 | 7.80 ±0.34* | 6.32±0.10* | 40.20±1.10* | 19.20±0.10* | 14.20 ± 1.56* |
| 5FU 25 | 10.24±0.81* | 4.22±0.10* | 50.20±0.30* | 17.22±0.23* | 21.33 ± 2.25* |
| 5FU 50 | 15.50±0.92* | 3.78±0.10* | 62.11±1.10* | 13.12±0.11* | 22.45 ± 2.56* |
| 5FU 100 | 21.12±0.91* | 2.52±0.11* | 82.20±1.5* | 11.14±0.04* | 24.21 ± 2.63* |

Mean ± S.E.M, \* $p < 0.05$ significant difference from control, number of rats per treatment group =10. 5FU: 5-Fluorouracil.

**Table 3:** Effect of 5-fluorouracil on cerebral cortex neurotransmitter levels.

| Groups | BDNF (pmol/mg tissue) | Acetylcholine (pmol/mg tissue) | Dopamine (pmol/mg tissue) | Serotonin (pmol/mg tissue) |
| --- | --- | --- | --- | --- |
| Control | 25.11±0.10 | 3.50±0.01 | 48.15±2.22 | 9.20±0.10 |
| 5FU 12.5 | 23.22±0.12 | 3.42±0.19 | 40.22±1.11* | 7.12±0.10* |
| 5FU 25 | 15.91±0.10* | 2.22±0.11* | 35.42±1.21* | 6.52±0.14* |
| 5FU 50 | 13.23±0.20* | 1.82±0.10* | 33.55±0.22** | 6.40±0.31* |
| 5FU 100 | 9.21±0.20* | 1.10±0.10* | 20.13±0.23* | 4.12±0.11<sup>#</sup> |

Data presented as Mean ± S.E.M, \*p < 0.05 vs. control, #p<0.05 significant difference from CYP, number of rats/group =6, BDNF: Brain derived neurotrophic factor.

### 3.7 Effect of 5-fluorouracil on neurotransmitter levels in the cerebral cortex

Table 2 shows the effect of 5-fluorouracil on neurotransmitter levels in the cerebral cortex. There was a significant decrease in the levels of brain derived neurotropic factor (BDNF) with 5FU at 12.5, 25, 50 and 100 mg/kg body weight compared to control. Levels of acetylcholine in the cerebral cortex also decreased significantly with 5FU at 12.5, 25, 50 and 100 mg/kg body weight compared to control Dopamine levels decreased significantly with 5FU at 12.5, 25, 50 and 100 mg/kg body weight compared to control. Serotonin levels also decreased significantly 5FU at 12.5, 25, 50 and 100 mg/kg body weight compared to control.

### 3.8 Effect of 5-fluorouracil on histomorphology of the cerebral cortex

Figures 6 (A, B, C, D and E) and 7 (A, B, C, D and E) are representative haematoxylin and eosin (H&E) and cresyl fast violet (CFV) stained sections of the rat cerebral cortex respectively. Examination of the H&E-stained slides of the group of rats administered vehicle revealed characteristic well delineated layers of the cerebral cortex. Scattered within the neuropil are multipolar shaped pyramidal cell with large vesicular nucleus, granule neurons with large open-faced nuclei and scanty cytoplasm and small sized neuroglial cells in rats administered vehicle (Figure 6A). The neuropil, which is pink staining in the H & E slides is well preserved in the CFV stained slides (Figure 7A). In the groups administered 5FU at 12.5 and 25 mg/kg body weight, normal shaped pyramidal cell can be observed interspersed between granule cells and neuroglial cells in the H&E Figure 6B and 6C) and CFV (Figure 7B and 7C) stained slides respectively. In the groups administered 5FU at 50 (Figure 6D and 7D) and 100 mg/kg (Figure 6E and 7E), observed were numerous neuroglial cells, degenerating pyramidal and granule cells interspersed between normal pyramidal and granule cells. Evidence of neuronal degeneration and neuronal injury included presence of pale-staining neurons with shrunken nuclei.

**Figure 6.**
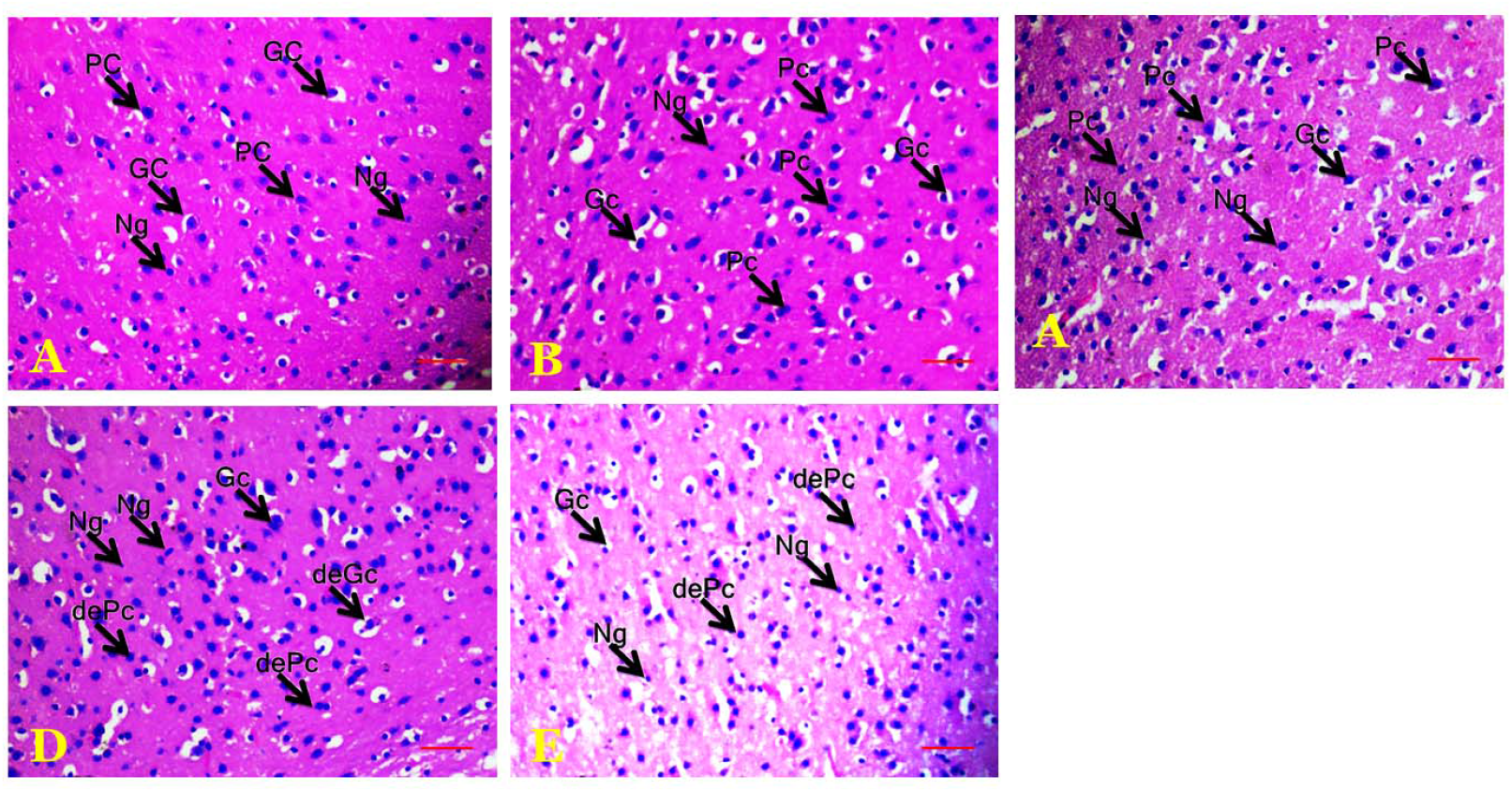
(A, B, C, D and E) are representative haematoxylin and eosin (H&E) stained sections of the rat cerebral cortex. PC: Pyramidal cells, GC: Granule Cells, Ng: Neuroglia, DePc: Degenerating Pyramidal cells. De Gc: Degenerating granule cells. Magnification: X100

**Figure 7.**
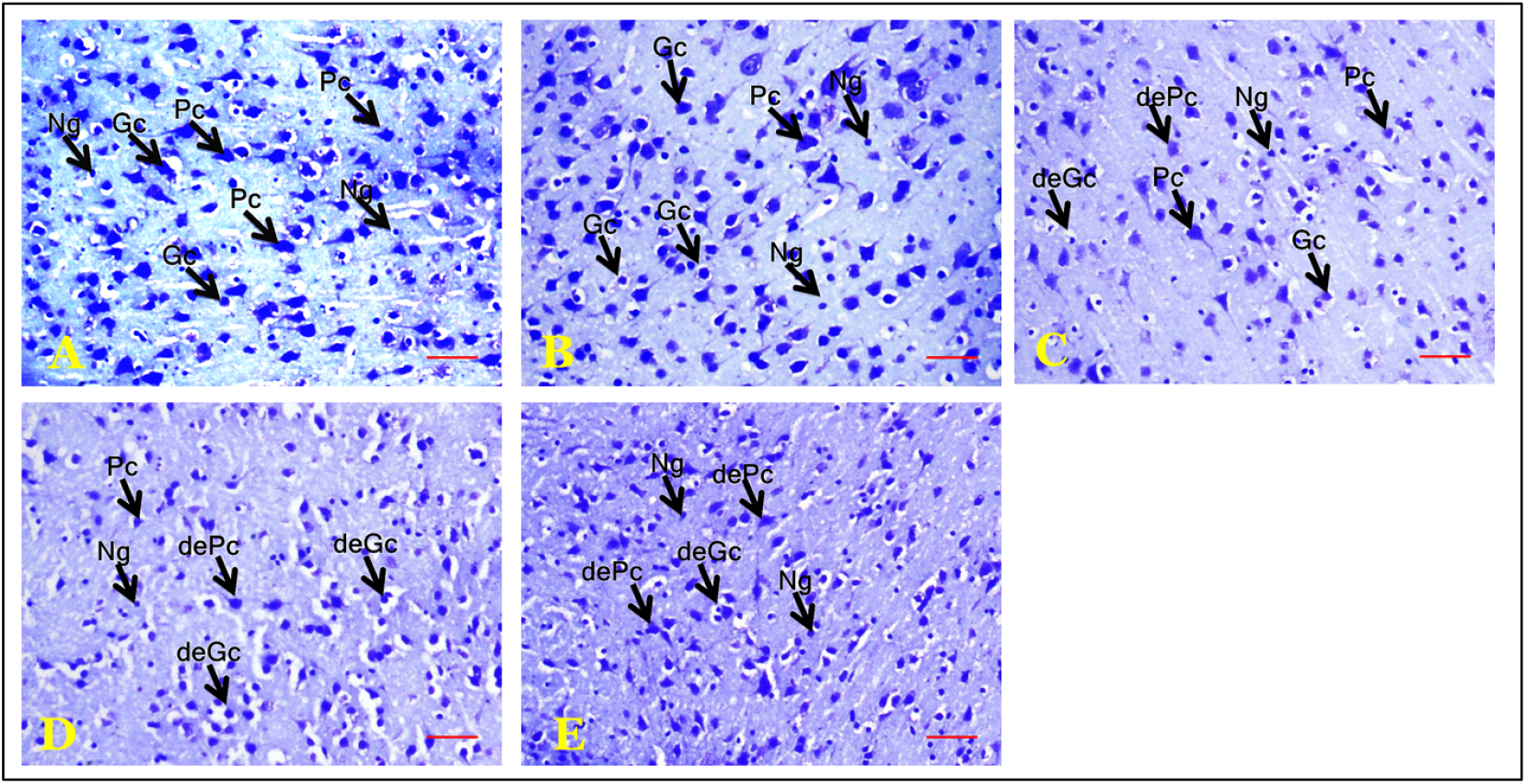
(A, B, C, D and E) are representative cresyl fast violet (CFV) stained sections of the rat cerebral cortex. PC: Pyramidal cells, GC: Granule Cells, Ng: Neuroglia, DePc: Degenerating Pyramidal cells. De Gc: Degenerating granule cells. Magnification: X100

### 3.9 Effect of 5-fluorouracil on Neuron Specific Enolase (NSE) immunoreactivity in the cerebral cortex

Figure (8 A-E) are representative NSE immunohistochemistry-stained sections of the rat cerebral cortex. Examination revealed the presence of numerous NSE-positive intact neurons with bluish staining nucleus and a brown rim in the groups administered vehicle (Figure 8A) and 5FU at 12.5 (Figure 8B) and 25 mg/kg (Figure 8C). The NSE-positive cells have pyramidal and granule cell morphologies, whilst the glial cells have no stain, appearing white. In the groups administered 5FU at 50 (Figure 8D) and 100 (Figure 8E) there is a decrease in the number of and intensity of NSE-positive cells. Also, the cell morphologies are altered with numerous degenerating pyramidal and granule cells.

**Figure 8.**
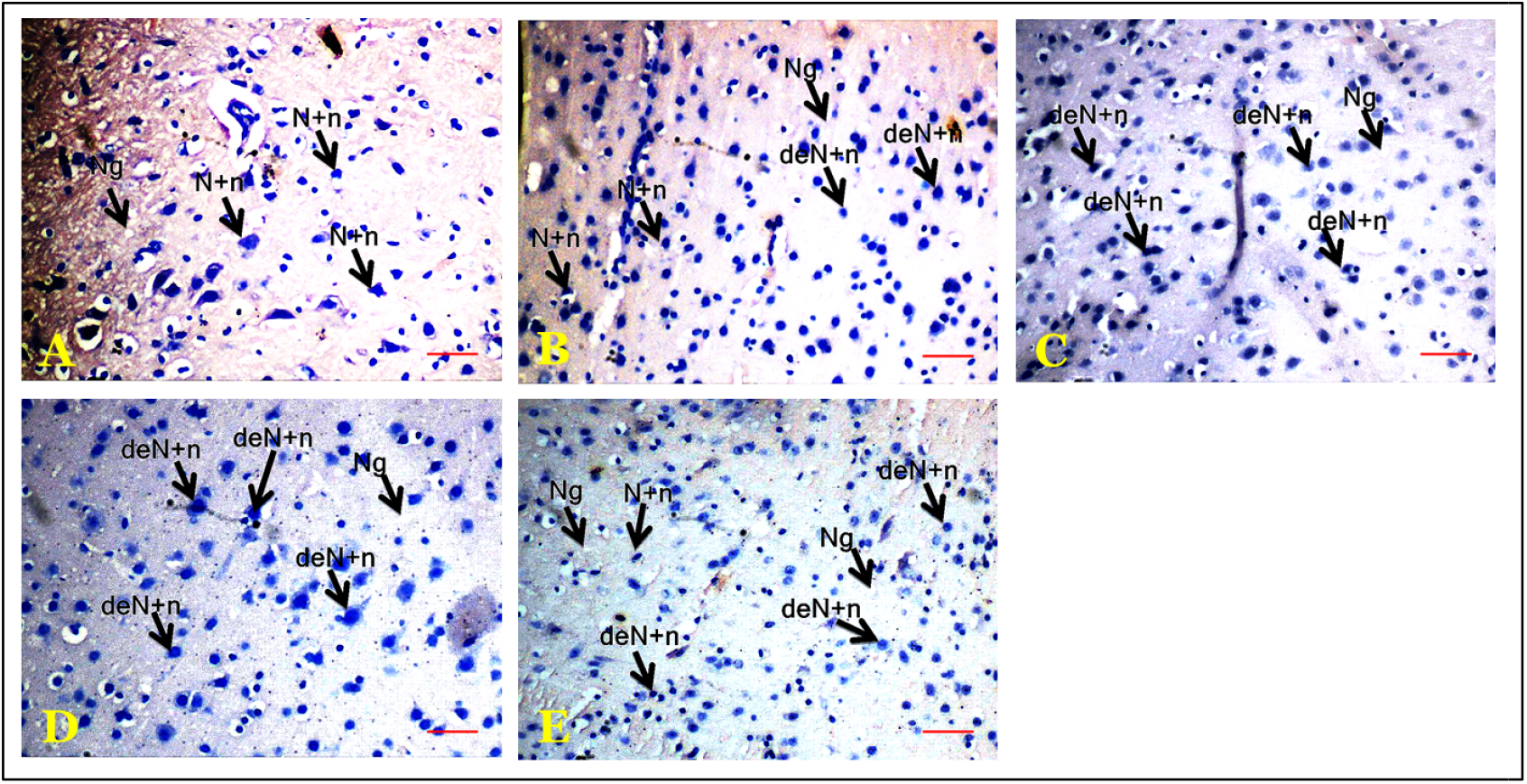
**(A-E)** are representative NSE immunohistochemistry-stained sections of the rat cerebral cortex. A: control, B5FU at 12.5 mg/kg body weight. C: 5FU at 25mg/kg body weight D: 5FU at 50 mg/kg body weight and E: 5FU at 100 mg/kg body weight. N+N neuron specific enolase reactive neurons, DeN+ n Degenerating Neuron specific enolase reactive neurons. Magx100

## Discussion

Advances in our understanding of the adverse nervous system effects of cancer chemotherapy agents would assist in determining regimens that considerably reduce the risk of neurotoxicity. Therefore, the current study evaluated the effects of four doses of 5FU on neurobehavioural, neurochemical, and neuromorphological changes in the cerebral cortex of rats. Results showed that 5-FU administration was associated with decrease in weight gain, feed intake, locomotor activity, rearing activity, self-grooming, spatial working memory, total antioxidant capacity, and interleukin 10 levels. Also observed were increased anxiety, lipid peroxidation and tumor necrosis factor– α and IL-6 levels. A decrease in the cerebral cortex levels of BDNF, dopamine, serotonin and acetylcholine levels were also observed.

In this study, the administration of 5-fluorouracil produced a marked reduction in both body weight and feed intake across all treatment groups when compared with controls. The decline in body weight is in line with a number of previous studies that had reported similar effects [36-38]; with feed intake and weight gain reduction. It also supports the well-documented systemic toxicity of 5-FU, which is known to affect rapidly dividing cells, including those of the gastrointestinal tract resulting commonly in mucositis, nausea, and gastrointestinal irritation reducing appetite and nutrient absorption, thereby contributing to weight loss [36-38]. The significant reduction in feed intake observed in treated animals likely reflects anorexigenic effects and gastrointestinal discomfort induced by 5-FU. Previous studies have reported similar findings, attributing decreased feed consumption to chemotherapy-induced alterations in taste perception, intestinal inflammation, and metabolic stress. Reduced nutrient intake further exacerbates weight loss, creating a vicious cycle of malnutrition and systemic weakening [39]. Also, the decrease in feed intake and resultant weight reduction observed following 5FU has also been linked to reduction in the MRNA encoding peptide YY [38]. Body weight and appetite loss in experimental animals are considered surrogate indicators of a generalized toxicity and declining health status [40]. The consistent decline across all 5-FU doses in this study reinforces the notion that even lower doses of the drug can impair feeding behavior and body mass regulation. These findings are in line with clinical observations in cancer patients receiving 5-FU, many of whom experience cachexia, anorexia, and weight loss during therapy [39].

Exploratory behaviours, measured as line crossing and rearing frequency, were significantly reduced in 5-FU–treated rats when compared with controls, with the decline being consistent across all dose groups. In the open field paradigm, line crossing reflects horizontal locomotor activity, while rearing is an index of vertical exploration and curiosity-driven behaviour [41]. Together, these measures provide insight into the animal’s general activity level, motivation, and emotional state. The results of this study are consistent with previous studies which had associated 5-FU administration with a significant reduction in the locomotor/exploration activity of C57Bl/6j mouse [42]. The reduction in line crossing observed across all dose groups in this study would suggests that 5-FU impairs spontaneous locomotor activity, which could be attributable to fatigue, muscle weakness, or central nervous system suppression of motor activity. Similarly, the marked reduction in rearing indicates diminished exploratory drive and environmental engagement, possibly reflecting reduced motivation, increased anxiety or both [41-43]. These behavioural alterations are in keeping with the concept of chemotherapy-induced sickness behaviour, characterized by lethargy, decreased activity, and loss of interest in the environment [43]. Clinically, these findings mirror seemingly patient reports of fatigue, psychomotor slowing, and reduced initiative during and after chemotherapy with fluoropyrimidines. Thus, the decreased line crossing and rearing observed in this study may represent preclinical correlates of chemotherapy-associated behavioural impairments.

Self-grooming is a complex, stereotyped behaviour in rodents that serves both a maintenance function (cleaning and thermoregulation) and an emotional function, as it is modulated by stress, anxiety, and central neurotransmitter activity. In the open field paradigm, changes in grooming frequency are often interpreted as indicators of the animal’s emotional reactivity, motivational state, and neurochemical balance [45-47]. In this study, 5-FU administration produced a significant reduction in grooming across all treated groups. This finding suggests that 5-FU exerts a suppressive effect on motivated, self-directed behaviours, which may reflect neurotoxicity, sickness behaviour, and emotional blunting [47]. Reduced grooming is consistent with chemotherapy-induced malaise and anorexia observed in parallel with body weight and feed intake reductions, pointing to a general decline in the animals’ well-being. Neurobiologically, grooming behaviour is strongly regulated by dopaminergic and serotonergic systems, particularly in the striatum, prefrontal cortex, and limbic structures [44, 48]. The central inhibitory response observe across all three behaviours could be associated with chemotherapy-induced oxidative stress and an increase in proinflammatory cytokines also observed in this study These changes could impair neural circuits underlying the initiation and sequencing of grooming patterns and behaviours generally. Additionally, inflammatory cytokines such as TNF-α and Il-6 are known to modulate motivational behaviour, and their upregulation may contribute to the observed suppression. A few studies have associated the increase in this cytokine with decreased appetite, fatigue, and other motivational suppression [49, 50].

In the elevated plus maze, 5-FU was associated with a significant reduction in the time spent in the open arms and a corresponding increase in the time spent in the closed arms across all doses. Since the open arms of the maze are perceived as aversive due to their height and exposure, while the closed arms provide relative safety, these results suggest that 5-FU enhances anxiety-like behaviour in rats; with higher doses evoking a higher anxiogenic response. This finding aligns with previous experimental reports demonstrating that systemic 5-FU administration increases indices of anxiety in rodents [42]. In the current study, the increased anxiety response may be related to the changes in the central neurotransmitters that were seen with 5-FU administration.

In this study, behavioural manifestations of the cognitive deficits associated with 5-FU administration were seen as reduced scores in the Y-maze and Radial arm mazes. The alternation index in the radial arm maze and the percentage alternation in the Y maze are established measures of spatial working memory, as they rely on the animal’s capacity to remember previously visited arms and avoid revisits within a short period [51, 52]. The observed impairments therefore indicate that 5-FU compromises short-term memory and spatial learning abilities. A previous report had shown spatial cognitive impairments following 5-FU administration [42]. 5-Fluorouracil administration was observed to evoke spatial cognitive impairments with the attendant morphological manifestation of a decreased neurogenesis within the hippocampus. Some other studies deduced that the cognitive impact of 5-FU can be linked to alteration of cell proliferation and neurogenesis in the dentate gyrus [53]. Also, a reduction in dendritic spines and increased in the inflammatory cytokines in the brain have been implicated [54]. The findings in our current study generally agree with these.

Biochemical analysis of the cerebral cortex revealed a clear shift toward oxidative stress and neuroinflammation following 5-FU treatment. Lipid peroxidation, indexed by malondialdehyde (MDA), increased significantly across all doses, indicating enhanced free radical generation and membrane lipid damage. This was accompanied by significant elevations in pro-inflammatory cytokines including tumour necrosis factor-α (TNF-α) and interleukin-6 (IL-6), while protective factors, including total antioxidant capacity (TAC) and the anti-inflammatory cytokine interleukin-10 (IL-10), were markedly reduced. This biochemical profile is consistent with the well-recognized role of 5-FU in generating reactive oxygen species (ROS) and disrupting redox homeostasis and is consistent with the results of other studies that had reported the association between 5-FU and the triggering of oxidation stress/lipid peroxidation [55, 56]. Excess ROS can trigger lipid peroxidation, protein oxidation, and DNA damage, thereby contributing to neuronal dysfunction [55, 56]. The parallel increase in pro-inflammatory cytokines suggests that oxidative imbalance may activate glial cells and perpetuate a cycle of neuroinflammation, which is known to impair synaptic plasticity and cognitive processing [57]. The reduction in IL-10, a critical anti-inflammatory cytokine, further underscores the loss of protective regulatory mechanisms, allowing unchecked inflammation to exacerbate neural injury [58, 59].

Administration of 5-FU was associated with a broad suppression of central cortical neurochemical systems critical for cognition, mood regulation, and behavioural control. Brain-derived neurotrophic factor (BDNF) levels decreased significantly across all treatment doses. BDNF plays a central role in neuronal survival, synaptic plasticity, and memory formation; its reduction suggests impaired neuroplastic capacity, which may underlie the observed deficits in spatial working memory and exploratory behaviours. Similarly, cortical acetylcholine levels were markedly reduced. Acetylcholine is a key neurotransmitter for attention, learning, and memory processes, particularly within prefrontal–hippocampal circuits. Its depletion provides a direct mechanistic link to the impaired alternation performance in both the radial arm and Y maze. In parallel, dopamine and serotonin levels were also significantly decreased by 5-FU. Dopamine regulates motivation, reward-driven behaviours, and motor activity, while serotonin modulates mood, anxiety, and affective states. The reductions in these monoamines align with the suppressed exploration, decreased grooming, and increased anxiety-like behaviour observed in the open field and elevated plus maze. Together, these changes indicate that 5-FU disrupts multiple neurotransmitter systems in a manner consistent with the emergence of apathy, psychomotor slowing, and heightened anxiety.

Histological evaluation of the rat cerebral cortex provided further evidence of 5-FU-induced neurotoxicity in a dose-dependent manner. In control animals, H&E and CFV-stained sections revealed a well-preserved cortical architecture, with clearly delineated cortical layers and normal populations of pyramidal neurons, granule neurons, and glial cells. Neurons displayed large vesicular nuclei and prominent nucleoli, consistent with healthy morphology, while the neuropil was intact and well stained. At low doses of 5-FU (12.5 and 25 mg/kg), cortical sections retained largely normal architecture, with preserved pyramidal and granule cells interspersed with glial cells. This suggests that at these doses, cortical neurons can partly resist or compensate for the cytotoxic effects of 5-FU. However, at higher doses (50 and 100 mg/kg), profound neuronal degeneration and architectural disruption became evident. Pyramidal and granule cells exhibited morphological hallmarks of neuronal injury, including pale-staining cytoplasm, shrunken or pyknotic nuclei, and loss of normal cell shape. These degenerative changes were accompanied by a marked increase in glial cell density, which may reflect gliosis, a reactive process secondary to neuronal injury. The CFV-stained sections further confirmed disruption of Nissl substance, indicative of impaired protein synthesis and neuronal metabolic stress.

Neuron-specific enolase (NSE) immunostaining further confirmed the dose-dependent neurotoxic effects of 5-FU on the cerebral cortex. Neuron-specific enolase is a neuronal subtype and one of the five isomers of the glycolytic enzyme enolase [60, 61]. It was initially identified in brain tissue extracts and later in the amine precursor uptake and decarboxylation (APUD) cells and neurons of the neuroendocrine system. In health it is localized within the central and peripheral neurons however it is not present in glial cells [60, 61]. In this study Immunohistochemical examination of the rat cerebral cortex revealed a decrease in the neuron specific enolase positive neurons with increasing doses of 5FU which is associated with neuronal injury. studies have reported.

## Conclusion

In conclusion these findings provide evidence that 5-FU induces dose-dependent neurobehavioural, neurochemical and neuronal degeneration in the cerebral cortex, with higher doses producing marked reductions in neuronal viability and functional integrity

## Notes

### Competing Interest Statement

The authors have declared no competing interest.

### Summary of Updates

The Title of the manuscript has been revised as have been some errors in the writing of the references

